# *De Novo* Growth of Methanogenic Granules Indicates a Biofilm Life-Cycle with Complex Ecology

**DOI:** 10.1101/667386

**Authors:** Anna Christine Trego, Evan Galvin, Conor Sweeney, Sinéad Dunning, Cillian Murphy, Simon Mills, Corine Nzeteu, Christopher Quince, Stephanie Connelly, Umer Zeeshan Ijaz, Gavin Collins

## Abstract

Methanogenic sludge granules are densely packed, small (diameter, approx. 0.5-2.0 mm) spherical biofilms found in anaerobic digesters used to treat industrial wastewaters, where they underpin efficient organic waste conversion and biogas production. A single digester contains millions of individual granules, each of which is a highly-organised biofilm comprised of a complex consortium of likely billions of cells from across thousands of species – but not all granules are identical. Whilst each granule theoretically houses representative microorganisms from all of the trophic groups implicated in the successive and interdependent reactions of the anaerobic digestion process, parallel granules function side-by-side in digesters to provide a ‘meta-organism’ of sorts. Granules from a full-scale bioreactor were size-separated into small, medium and large granules. Laboratory-scale bioreactors were operated using only small (0.6–1 mm), medium (1–1.4 mm) or large (1.4–1.8 mm) granules, or unfractionated (naturally distributed) sludge. After >50 days of operation, the granule size distribution in each of the small, medium and large bioreactor types had diversified beyond – to both bigger and smaller than – the size fraction used for inoculation. ‘New’ granules were analysed by studying community structure based on high-throughput 16S rRNA gene sequencing. *Methanobacterium*, *Aminobacterium*, *Propionibacteriaceae* and *Desulfovibrio* represented the majority of the community in new granules. H2-using, and not acetoclastic, methanogens appeared more important, and were associated with abundant syntrophic bacteria. Multivariate integration analyses identified distinct discriminant taxa responsible for shaping the microbial communities in different-sized granules, and along with alpha diversity data, indicated a possible biofilm life cycle.

**Importance:** Methanogenic granules are spherical biofilms found in the built environment, where despite their importance for anaerobic digestion of wastewater in bioreactors, little is understood about the fate of granules across their entire life. Information on exactly how, and at what rates, methanogenic granules develop will be important for more precise and innovative management of environmental biotechnologies. Microbial aggregates also spark interest as subjects in which to study fundamental concepts from microbial ecology, including immigration and species sorting affecting the assembly of microbial communities. This experiment is the first, of which we are aware, to compartmentalise methanogenic granules into discrete, size-resolved fractions, which were then used to separately start up bioreactors to investigate the granule life cycle. The evidence, and extent, of *de novo* granule growth, and the identification of key microorganisms shaping new granules at different life-cycle stages, is important for environmental engineering and microbial ecology.

## INTRODUCTION

Biofilms form in a wide range of natural and built environments, and have important significance for biogeochemical cycling in Nature, as well as for clinical and industrial applications. Moreover, evidence suggests that most microorganisms form, or can be found in, complex biofilm aggregates (1). Aggregation is an ancient process that has allowed prokaryotic life to thrive even in the harshest of environments (2). However, though biofilms are classically found as layers, or films, attached to suitable surfaces – from rocks, to medical devices, to ship hulls – aggregation may also occur due to self-immobilisation of cells into discrete structures, such as flocs or granules, without the involvement of a surface. Many such examples can be found in engineered environments, such as in biological wastewater treatment, where prevailing conditions of shear, and hydrodynamic, stresses promote flocculation and granulation. Common types include anaerobic ammonium oxidising (annamox) granules (3), aerobic granules (4), and anaerobic (methanogenic) granules (5).

Indeed, the success of high-rate anaerobic digestion (AD) – which is widely applied to treat a range of industrial wastewaters – is underpinned by the spontaneous generation of active biomass in the form of anaerobic granules (AnGs) (Fig 1), which are small (approx. 0.5-2.0 mm), densely-packed biofilm spheres comprising the complex microbial community necessary for the complete mineralisation of organic pollutants by AD (6). The settleabilty of AnGs accounts for long biomass retention – even in ‘upflow’ bioreactors, such as the upflow anaerobic sludge bed (UASB) and expanded granular sludge bed (EGSB) bioreactors, operated with short hydraulic retention times (HRT), and very high volumetric loading and up-flow velocities (7). A single granule can contain billions of microbial cells from thousands of species juxtaposed, and immobilised, within a complex matrix of extracellular polymeric substances (EPS). Within these highly organised consortia, a collection of microbial trophic groups mediates a cascade of interdependent reactions resulting in complete degradation of complex organic wastewater pollutants. Equally, the consortium’s species rely on efficient mass transfer of substrates into, and throughout, the granule.

**Figure 1.**
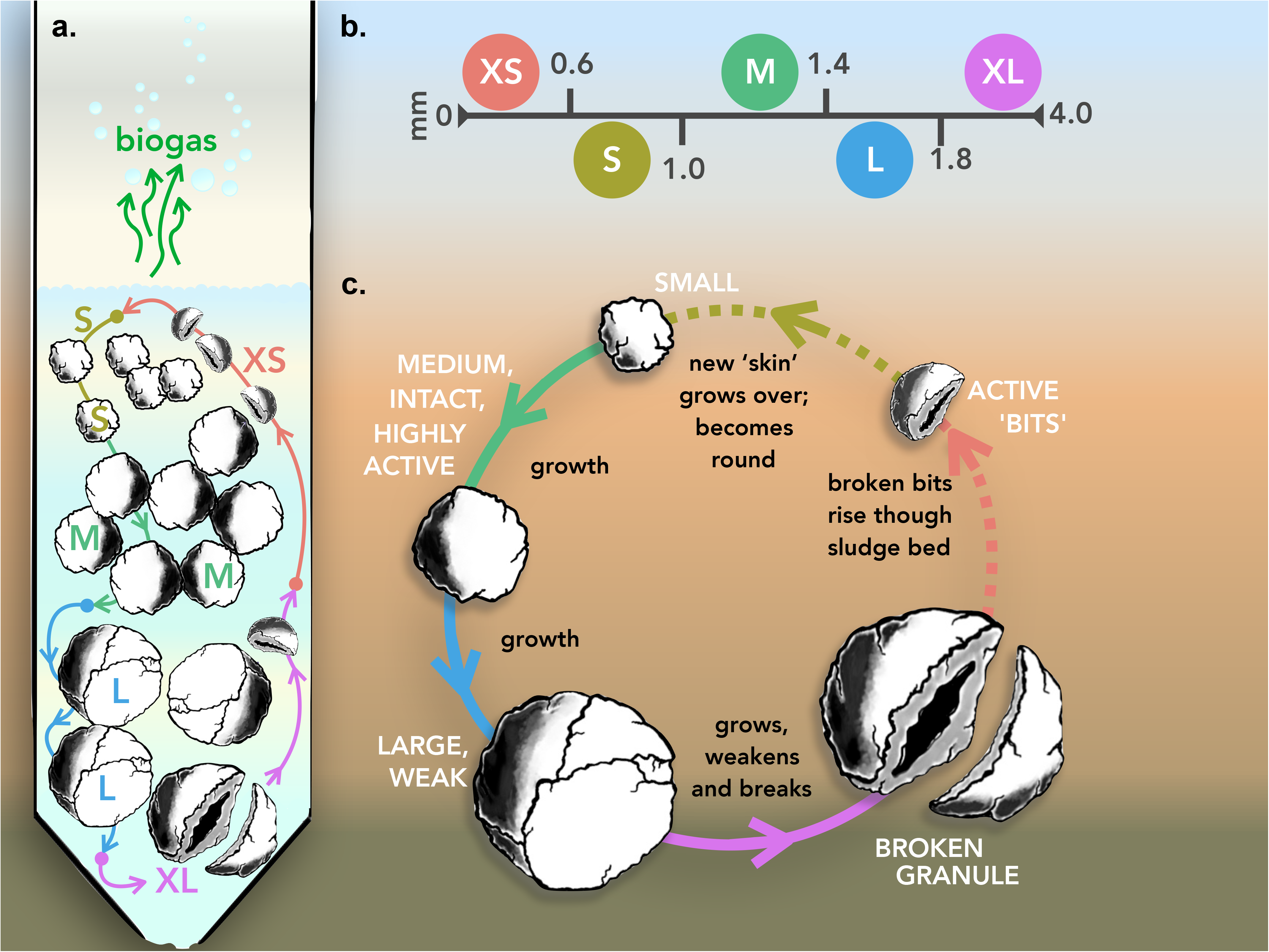
Granular growth hypothesis and biofilm life-cycle model. **(a)** operation of the model inside an anaerobic bioreactor; **(b)** size fraction parameters; and **(c)** the generalised growth model.

Granulation is a process whereby suspended particles and planktonic cells accumulate, forming small dense biofilm aggregates (6). Unlike conventional biofilm formation, which is a well-documented phenomenon, less is understood about granulation of anaerobic sludge. Hulshoff Pol et al. (8) comprehensively reviewed the topic, summarising the various theories proposed on granulation, which they categorised as physical, microbial or thermodynamic. However, none has been solely accepted as a ‘unified theory on anaerobic granulation’ (9). The only consensus seems to be that *Methanosaeta concilii*, an acetoclastic methanogen, is a key organism in the process (8) due to its filamentous morphology. These archaea can either (i) aggregate together, (ii) attach to suspended particles, or (iii) potentially form a bridge between existing microflocs – aiding in the critical first step of forming granule precursors (10–12). Moreover, whilst several theories have been promoted on granulation mechanisms, the growth, development and evolution of AnGs has been largely overlooked.

An intensive characterisation of anaerobic granules by (13) produced a granular growth hypothesis and life-cycle model, built upon pragmatic observations of previous studies suggesting small granules are ‘young’ and large granules are ‘old’ or mature (14–16). The granular growth hypothesis proposed small granules grow by cell accumulation and replication – which is similar to classical biofilm growth models (2, 17) – into medium-sized, intact, and highly-active granules. These grow into large granules, but the structure weakens as external shear stresses, and gas diffusion from the interior of the biofilm, cause cracks and voids in the biofilm structure (16). The hypothesis proposes the fate of the largest, oldest granules, as breaking apart into smaller biofilm ‘bits’ or layers. The broken parts, however, are still comprised of active biomass and eventually round over to form the basis for new, small granules – the entire process being cyclical (Fig 1). Such a model for granule growth along a predictable life-cycle inside anaerobic bioreactors could not only provide opportunities for precision management of sustainable, efficient wastewater treatment applications, but also improve our understanding of microbial community assembly and succession in dynamic biofilms.

Many studies have focused on granulation (8), and associated dynamics of physico-chemical properties and microbial community structure. Fewer studies, however, have attempted to follow the fate of granular biofilms over their entire life. Indeed, microbial aggregates provide potentially fascinating opportunities as “parallel evolutionary incubators”, as previously suggested (18). Intense interest now surrounds the study of granular biofilms as playgrounds to investigate fundamental concepts in microbial ecology (19). Recent studies (20, 21) have used granules – albeit aerobic, and not anaerobic, granules – to study the roles of phenomena, such as immigration and species sorting, driving microbial community assembly.

The purpose of this study was to test the granular growth and life-cycle hypothesis; to determine whether granules do, indeed, grow and develop in a predictable way, from small to medium and, finally, to large. This is the first experiment, of which we are aware, to compartmentalise granules into size-resolved fractions (small, medium and large), which were then used to separately start up bioreactors to investigate the granule life cycle. Moreover, undisturbed sludge, providing a ‘meta-community’ and full complement of size fractions, was used as a comparator. The extent, nature and ecology of ‘new’ granules emerging in the experiments was monitored.

## RESULTS

### Bioreactor performance

The twelve laboratory-scale bioreactors set up across the four conditions tested – i.e. R_S1_-R_S3_ containing only ‘S’ granules (Ø, 0.6-1.0 mm), R_M1_-R_M3_ containing only ‘M’ granules (1.0-1.4 mm), R_L1_-R_L3_ containing only ‘L’ granules (1.4-1.8 mm), and R_N1_-R_N3_ containing a full complement of unfractionated, naturally distributed (N) sludge (S, M and L, as well as XS and XL) – allowed the emergence of new granules to be detected and studied. The operating conditions were selected to mimic, as closely as possible, the conditions of the original full-scale bioreactor with respect to mean temperature, upflow velocity, organic loading rate and feed type (Table 1). Each of the three sets of bioreactors performed similarly. During the first week (Phase 1), influent pH decreased (mean, pH 4.1) in each of the bioreactors. After supplementation of the influent with sodium bicarbonate, the pH stabilised (mean, pH 7.8) over the remainder of the experiment. Biogas methane concentrations were low during the initial acidification experienced during Phase 1 but increased during Phase 2 (Fig 2).

**Table 1.** Full- and laboratory-scale bioreactor operating parameters.

| **Parameter** | **Full-scale bioreactor** | **Laboratory-scale** |
| --- | --- | --- |
| Influent pH | 6.3 | 3.9 – 9.5 |
| Operating Temperature | 37°C | 37°C |
| Upflow velocity | 1.2 m h<sup>-1</sup> | 1.2 m h<sup>-1</sup> |
| Influent COD Concentration | 4.5 g L<sup>-1</sup> | 15.7 g L<sup>-1</sup> |
| Organic loading rate | 15.7 g COD L<sup>-1</sup> d<sup>-1</sup> | ~15.7 g COD L<sup>-1</sup> d<sup>-1</sup> |
| Hydraulic retention time | 6.86 h | 24 h |
| Wastewater type | Potato-processing | Synthetic (Table 2) |

**Figure 2.**
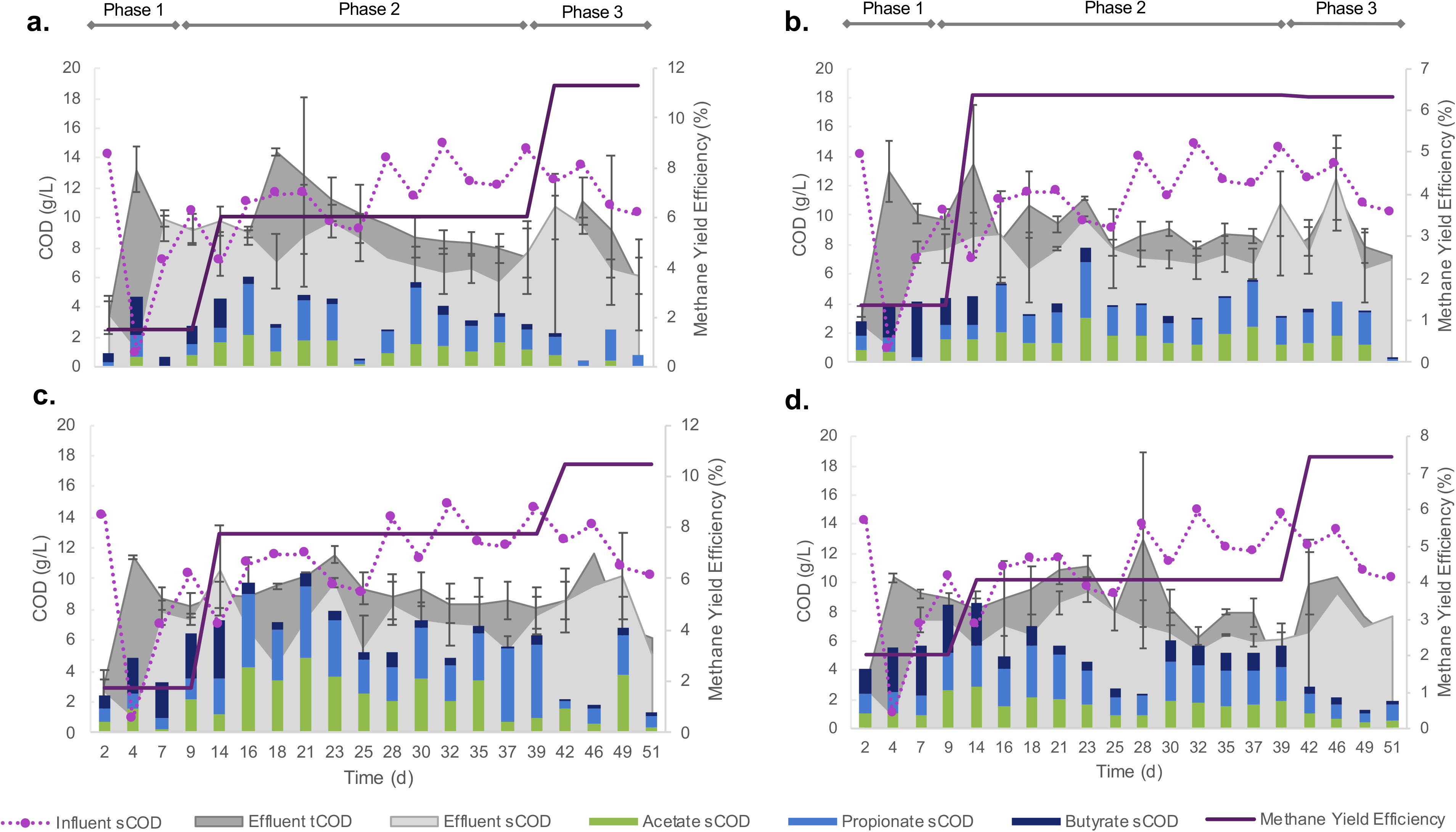
Methane yield efficiency; COD conversions; and key VFA (acetate, propionate and butyrate) contributions to effluent sCOD in each of the four bioreactor sets: **(a)** R_S1_ – R_S3_; **(b)** R_M1_ – R_M3_; **(c)** R_L1_ – R_L3_; **(d)** R_N1_ – R_N3_.

Measurements of total chemical oxygen demand (tCOD) include both soluble (sCOD) and particulate COD (pCOD), and so the difference between tCOD and sCOD measurements indicates the concentration of pCOD. During Phase 1, more COD left the bioreactors than was fed to them (Fig 2). However, only soluble COD (and no pCOD) was fed to the bioreactors (sCOD was equal to tCOD in influent) and most of the tCOD in effluent during Phase 1 appeared as pCOD (Fig 2), indicating the COD mostly reflected sludge washout during the first week. This was largely reversed for the remainder of the trial, and COD removal improved significantly over the subsequent approximately four weeks (Phase 2), culminating in roughly 50% sCOD removal efficiencies by each of the bioreactors. Nonetheless, COD removal was lower again during the final approximately two weeks of the trial (Phase 3; Fig 2). Acetate, propionate and butyrate contributed to 50-90% of effluent sCOD (Fig 2).

Biomass washout was observed from each bioreactor variously over the course of the 51-d experiment, including from the ‘naturally-distributed’ condition (R_N1-3_). Bioreactor R_N2_ failed – and was stopped – on day 22, due to the loss of 52% of the sludge. The remaining 11 bioreactors experienced losses reaching up to 50%. Washout of biomass was noted throughout the trial, but increased during Phase 3 (as evidenced by higher pCOD concentrations; Fig 2). A net gain in biomass was observed in only two bioreactors, R_L1_ and R_L3_ (Fig S1).

### Shifts in granule size distribution

Size fractionation of biomass at the conclusion of the trial showed that the distribution of granule sizes had changed, and new granules – or ‘emerging sizes’ – were apparent in all of the bioreactors (Fig 3). Whereas granules were initially only one size, many new sizes had emerged after 51 days. In all three of the R_L_ bioreactors, and in two of the R_M_ bioreactors, a full range of sizes (from the XS, S, M, L, XL classifications) had emerged (Fig 3). In the two surviving R_N_ bioreactors, granules each of the five size classifications were still present, although the proportion of granules in M or above had increased. In fact, with only the exception of L granules in R_S2_ and R_M2_, and XL granules in R_S2_ & R_S3_, all five sizes emerged from all bioreactors (Fig 3).

**Figure 3.**
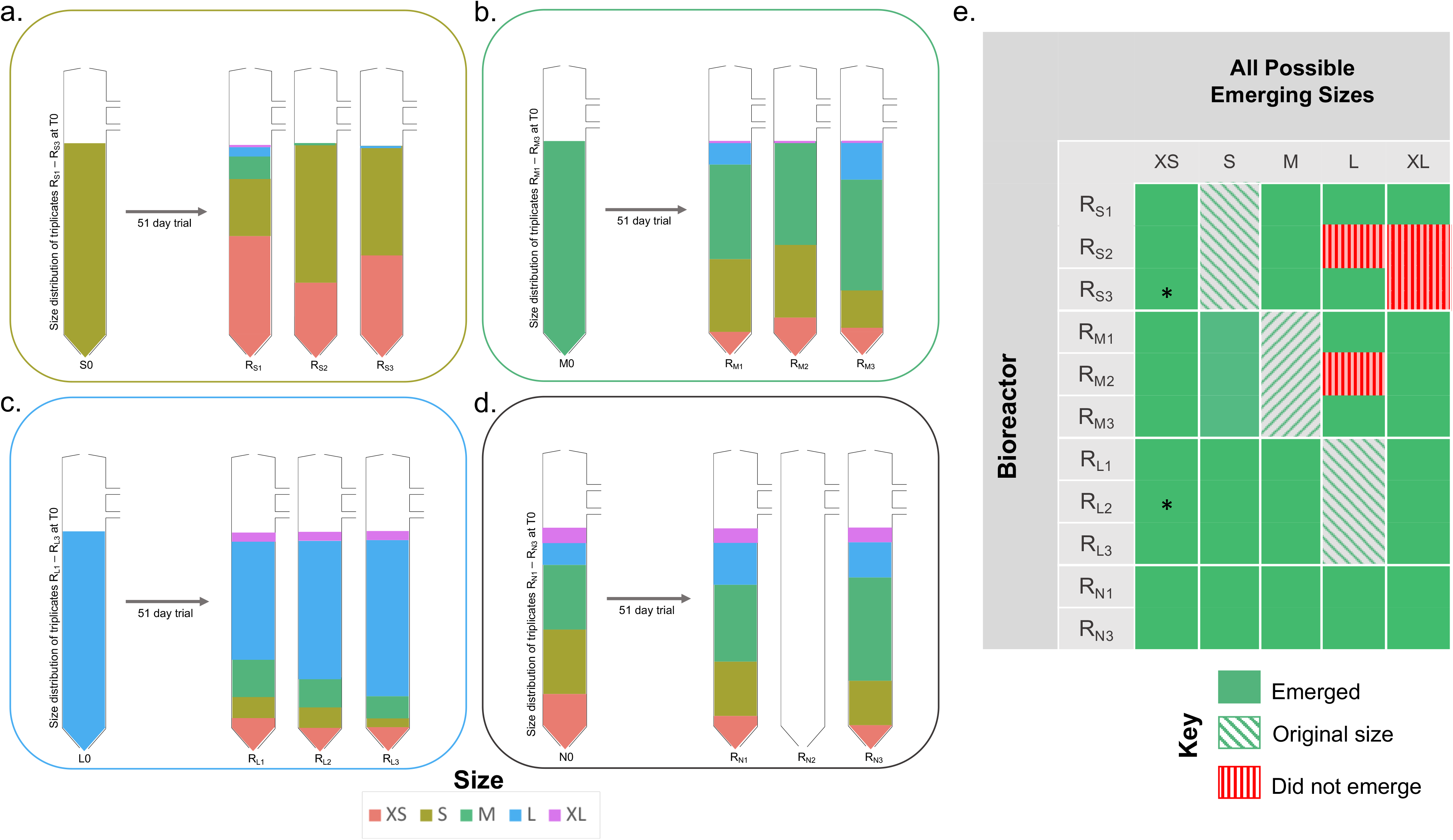
Changes in distribution of granules sizes in the R_S_, R_M_, R_L_ and R_N_ bioreactors during the trial (day 0 and each of the respective bioreactors at day 51), showing: **(a)** R_S1_ – R_S3_; **(b)** R_M1_ – R_M3_; **(c)** R_L1_ – R_L3_; **(d)** R_N1_ and R_N3_ bioreactors. Colours indicate the size of the emerging granules and their proportion of the total biomass present. **(e)** Map indicating frequency of observations of emerging sizes across the experiment.. No sequencing data available for samples marked with (*).

### Microbial community structure of emerging granules

Alpha diversity measurements, using Shannon Entropy, indicated similar trends for emerging granules from the R_M_ and R_L_ bioreactors (Fig 4). A linear reduction in alpha diversity – similar to the trend previously observed (13)– was apparent from S through to XL granules (i.e. there was more diversity in the microbial communities found in S granules than in bigger ones). Nonetheless, the alpha diversity in XS granules was significantly lower than in S granules – rather than *higher* as might have been expected based on previous findings (13). In fact, the diversity found in XS granules was similar to the diversity in XL granules (Fig 4).

**Figure 4.**
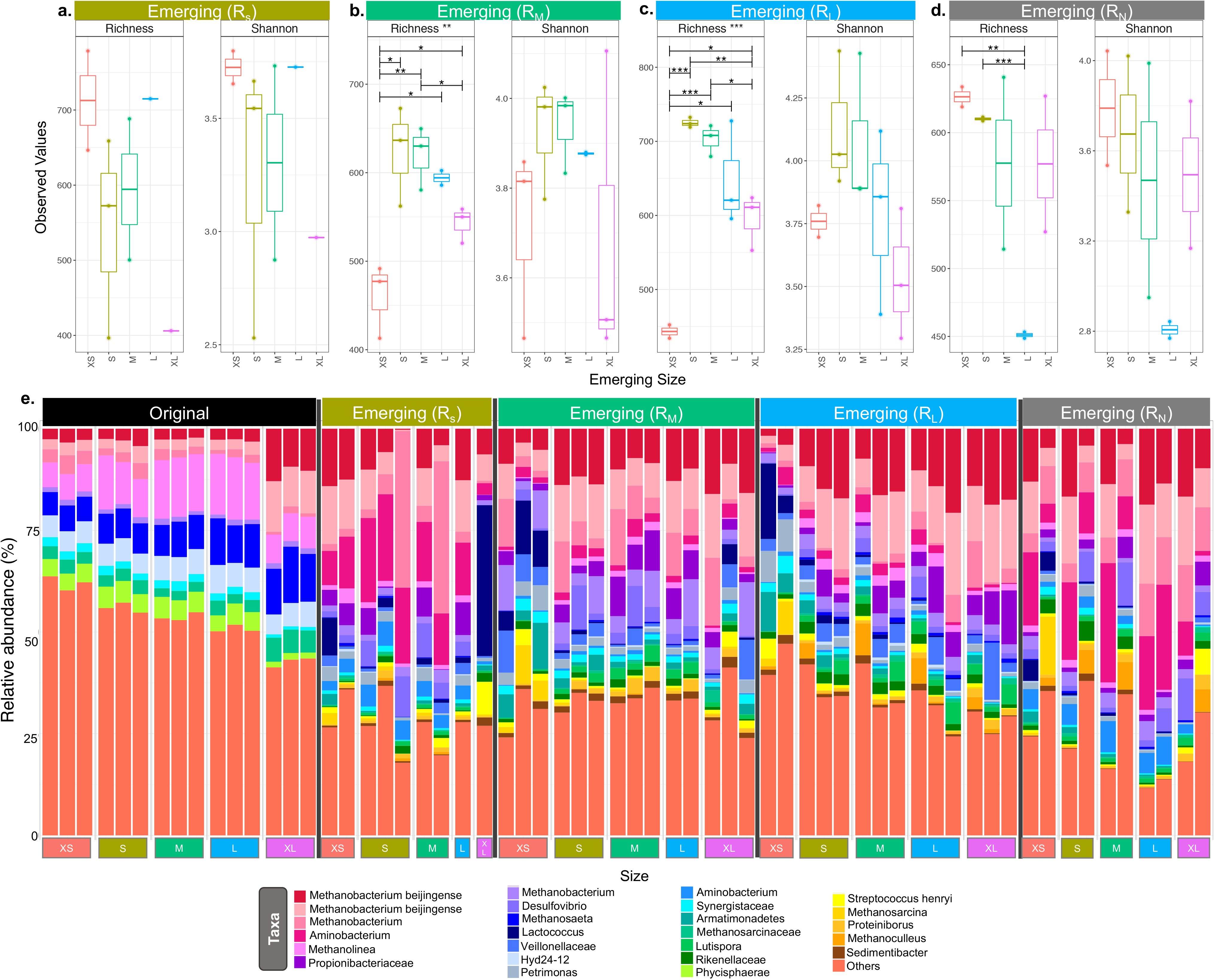
Box plots **(a-d)** of rarefied richness of ‘emerging sizes’ from across the four bioreactor sets: **(a)** R_S1_ – R_S3_; **(b)** R_M1_ – R_M3_; **(c)** R_L1_ – R_L3_; **(d)** R_N1_ and R_N3_; and bar chart **(e)** showing the top 25 relatively most abundant OTUs in original and new granules.

The initial (Day 0) community structure comprised of a mix of hydrogenotrophic (*Methanobacterium*, *Methanolinea*) and acetoclastic (*Methanosaeta*) methanogens (archaea). At the same time, the bacteria found to be relatively most abundant were generally all heterotrophic fermenters. Over the course of the trial, the make-up of the most abundant taxa shifted considerably. Across all of the new (or growing) granules – i.e. the emerging sizes from the bioreactors – the community structure was dominated by four operational taxonomic unit (OTU) classifications of *Methanobacterium*, in many cases accounting for 25-50% of the relative abundance of all taxa (Fig 4). Interestingly, *Methanosaeta* completely disappeared from amongst the 25 most abundant OTUs. Other highly abundant taxa included *Aminobacterium*, *Propionibacteiraceae* and *Desulfovibrio*.

Multivariate integration (‘MINT’) algorithms used for study-wise discriminant analyses (see Supplemental Material) identified a total of 38 ‘discriminant’ OTUs from two selected ‘components’ (Fig S2). Discriminant OTUs formed two phylogenetic clades from 11 distinct phyla. Mean relative abundances of these OTUs showed two general groupings: (i) those OTUs more abundant in either, or both, of the emerging XS and XL sized granules, and (ii) those OTUs which were more abundant in the emerging S, M, and L granule sizes.

## DISCUSSION

### Emerging sizes: granules grow

This study demonstrates that methanogenic granules in anaerobic digesters do, indeed, ‘grow’. In each of the nine bioreactors started up with granules from a discrete size classification (Fig 5), the final distribution of granule sizes shifted to include new (or ‘grown’) granules that were either larger or smaller than the original granules (Fig 3, Fig). The emergence of larger granules almost certainly indicates the growth of granules due to cell replication and the accumulation of formerly planktonic cells from the surrounding environment. The emergence of granules smaller than the original biomass might be explained in two ways: that (i) completely new granules formed from planktonic cells in the wastewater and the granulation process was continually initiated inside the digester, or (ii) bits and pieces of older, larger granules broke away and provided the foundation for new, small granules. The second explanation also points to a potential life-cycle of methanogenic granulation. What is actually likely, we suggest, is that both phenomena proceed simultaneously.

**Figure 5.**
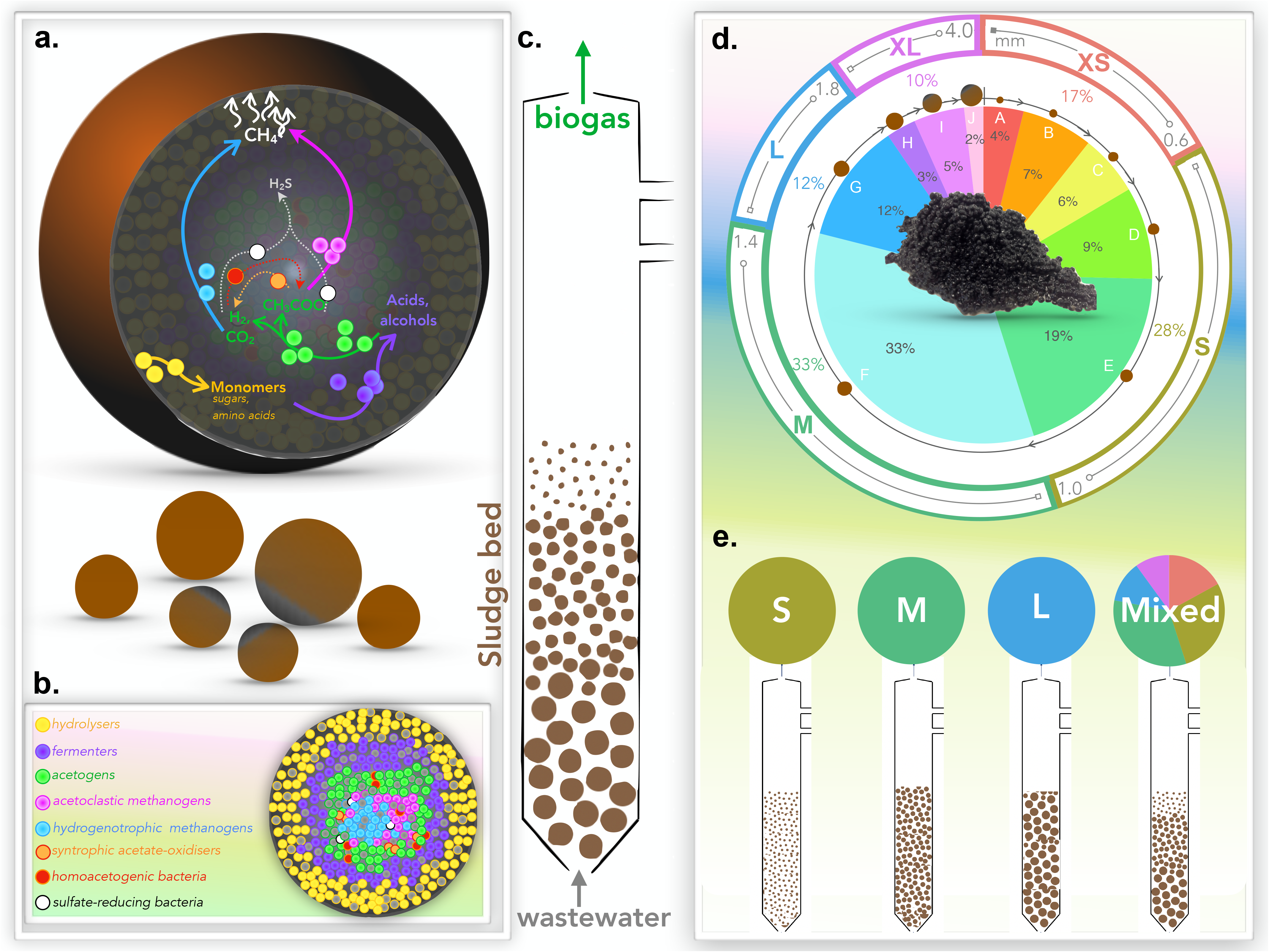
Schematics illustrating: **(a)** the AD pathway of organic matter degradation in the context of a granule; **(b)** theoretical distribution of the main trophic groups catalysing the process; **(c)** the engineered bioreactor system used to apply granules for wastewater treatment and biogas generation; **(d)** size distribution of biomass whereby the ten size fractions used by (Trego et al., 2018) were binned for this study into five size groups: extra-small (XS), small (S), medium (M), large (L), and extra-large (XL); and **(e)** the experimental set-up used to test granular growth where bioreactors were inoculated with either S, M, L or the naturally distributed (mixed) biomass.

An important component of the experiment was the set of bioreactors (R_N_) started up with a full complement of granule sizes, representing a sort of ‘meta-community’ of individual ecosystems (individual granules) – inspired in part by the recent description (18) of soil aggregates as parallel incubators of evolution. In the R_N_ bioreactors, the size distribution shifted during the experiment toward larger granules. This could be due to growth, and/or the operational conditions of the bioreactors selecting for larger sizes (i.e. the hydraulic regime and shear stresses applied). Another possible explanation could be that the sludge lost from the R_N_ bioreactors during the experiment included smaller granules although there was no indication that smaller granules were preferentially lost from any of the other (R_S_, R_M_ or R_L_) bioreactors. Indeed, for example, many XS granules, which emerged in the R_S_ bioreactors, appeared to resist washout and were retained in those bioreactors.

### Emerging pathways: dominance of hydrogenotrophic methanogenesis

The predominant members of the emerging microbiome across all of the samples included *Methanobacterium*, *Aminobacterium*, *Propionibacteriaceae* and *Desulfovibrio* species. Hori et al. (22) that found low pH and increasing VFA concentrations in anaerobic digesters resulted in more abundant *Methanosarcina* (acetoclastic & hydrogenotrophic methanogens) and *Methanothermobacter* (hydrogenotrophic methanogens) but fewer *Methanoculleus* (also hydrogenotrophic), concluding that VFA accumulation strongly influences archaeal community structure. Kotsyurbenko et al. (23) subsequently expounded this generalised conclusion, finding temporally falling pH in an acid peat bog shifted community structure from acetoclastic to hydrogenotrophic methanogens, concluding that pH shapes methanogenic pathways.

This was also supported by our experiment. *Methanosaeta* – an acetoclastic methanogen, which was abundant in the granules on Day 0 – was not detected in the emerging granules, whilst *Methanobacterium* – autotrophic, H_2_-using methanogens (24–28) also capable of formate reduction (29) – were dominant and likely feeding on increased dissolved hydrogen resulting from the accumulating VFA (30). *Propionibacteriaceae* – a family of heterotrophic glucose fermenters, producing propionate and acetate as primary products (31) – were also abundant in new granules, likely as VFA-producing acetogens. It is, of course, interesting to observe that granules emerged in this experiment without the apparent dominant involvement of the filamentous *Methanosaeta* spp., which tends to contradict the conventional understanding of granulation microbiology.

### Emerging ecology: supporting syntrophic relationships

The dominance of hydrogenotrophic methanogens (Fig S3) in the emerging granules appeared to support the abundance of syntrophic bacteria, including *Aminobacterium* – heterotrophic fermenters of amino acids that grow well with methanogenic, H_2_-consuming partners, such as *Methanobacterium* (32, 33) – and *Desulfovibrio* – sulfate-reducing bacteria (SRB) widespread in the environment (34), where they respire hydrogen or organic acids (35) often in syntrophy with methanogens (36). Interspecies metabolite exchange and hydrogen transfer (37) between syntrophic partners is critical in AD because the oxidation of organic acids and alcohols by acetogens may be thermodynamically feasible only when hydrogenotrophic methanogens (in this case, likely the *Methanobacterium*) consume, and maintain sufficiently low concentrations of, H_2_. It is clear that the microbial community – including in the emerging granules – responded to the prevailing environmental conditions within the bioreactors. Indeed, had there not been an accumulation of VFA in the bioreactors and a striking dominance of the H2-oxidising methanogens, a different community – perhaps characterised more strongly by the acetoclastic methanogens, such as *Methanosaeta* – may have developed.

### Emerging discriminants: size-specific OTUs

In general, the communities of all emerging granules were very similar with some, though few, significant differences in alpha diversity and rarefied richness. Nonetheless, 32 study-wise discriminants could be identified, using MINT-sPLS analysis, which were responsible for minor community shifts across the emerging granules from each bioreactor set. Phylogenetically, these discriminants formed two distinct clades – the first made up primarily of the phyla Firmicutes, Synergistetes and Chloroflexi, and the second clade comprising of Proteobacteria, Spirochaetae, Bacteroidetes, and Euryarchaeota. Many of the discriminant OTUs were generally upregulated in the emerging S, M or L granules, or were upregulated in either or both XS and XL granules. For example, *Lactococcus*, a glucose fermenter and primary member of the lactic acid bacteria group, and *Stenotrophomonas*, a likely nitrate reducer, were both upregulated in XS and XL granules, but rare in emerging S, M and L granules. Conversely, other taxa, such as the *Phycisphaerae*, *Leptospiraceae* and *Bdellovibrio*, were upregulated in the emerging S, M and L granules but infrequent in XS or XL granules.

Rather than observing a linear trajectory in diversity – from the smallest toward the largest granules – and a clear grouping of discriminants according to granule size (13), a more puzzling pattern manifested from this study. Coupling the patterns followed by the discriminant OTUs with patterns in alpha diversity, the microbial communities of XS and XL granules appeared to be more similar than previously observed (Fig 4) – thus pointing towards closing the loop on a life-cycle model for granular biofilms.

### Granular growth hypothesis and biofilm life-cycle

The granular growth hypothesis and biofilm life-cycle model proposes that granules start small, and through cell replication and biomass accumulation, swell into medium and then large aggregates. However, it postulates, based on previous evidence (16), that the larger the granule becomes, the more structurally unstable it is, and that it eventually breaks apart. These broken bits, still containing an active microbial community eventually round off (due to shear forces within the digester) and become the basis for new, small granules, so that the process is cyclical (Fig 1). The main objective of this experiment was to arrive closer to determining whether a life-cycle applies to methanogenic granules.

To accept the granular growth hypothesis we would need to see that bioreactors initially containing only small granules, would eventually contain medium, then large and, finally, extra-large granules. An equivalent scenario would be observed for each bioreactor set. Equally, clear trends in microbial community structure might be observed across the different sizes. For example, an XL granule would have a similar community structure to an XS granule, but may be significantly different to an S or M granule.

This study provides evidence for ‘growing’ granules and for the emergence of *de novo* granules. Granule growth was apparent in all nine of the R_S_, R_M_ and R_L_ bioreactors. Indeed, most contained granules – albeit, sometimes very few – from each of the five size classifications used. What remains unclear is the rate at which this happened, the mechanisms driving this process, and whether the process really is cyclical. For example, even if granules do break apart to form smaller, ‘new’ ones, whether there is a critical point (e.g. size or age) at which this happens is unresolved. This study would suggest, based on emergence of XS granules in the R_S_ bioreactors (Fig 3), that even small granules can break apart.

Analyses of the microbiomes of the emerging granules found, in some instances, a cyclical pattern in which the alpha diversity of XS and XL granules was similar (Fig 4). However, likely due to bioreactor operation, which shifted the community structure across all of the experiments, it remains unclear how the microbiome of anaerobic granules changes as they grow.

Meanwhile, although this experimental design provides an interesting perspective and means to uncover the trajectory and fate of granular biofilms, each size-controlled set of bioreactors started with a different, and constrained, microbial consortium. Thus, emerging granules from different bioreactor set-ups, although perhaps similarly sized, are not necessarily comparable.

In summary, granules were demonstrated to be dynamic aggregates inside anaerobic digesters, appearing to follow a progressive growth pattern from S, to M to L. XS granules emerged in all bioreactors, regardless of the starting size distribution. These either formed *de novo*, from the aggregation of free cells, or as a result of larger granules breaking apart. Further experiments should be done, under more stable bioreactor conditions, and with more intensive sampling regimes, to provide more evidence. The results of experiments based on innovative approaches to track the fate of growing granules will provide invaluable information to environmental engineers running bioreactors and to microbial ecologists studying community assembly phenomena, alike.

## MATERIALS AND METHODS

### Source and fractionation of biomass

Anaerobic sludge was obtained from a full-scale (8,256 m^3^), mesophilic (37ºC), EGSB bioreactor, inoculated with sludge granules from the Netherlands and treating potato-processing wastewater, in Lurgan, Northern Ireland. The full-scale bioreactor was operated at an upflow velocity of 1.2 m h^−1^ and an HRT of 6.86 h.

A comprehensive analysis of the granules across a highly resolved size distribution was previously performed (13). The ten size fractions (A-J) characterised in that study were grouped, for this study (Fig 5), into five distinct size classifications: extra-small (XS), small (S), medium (M), large (L), and extra-large (XL). Granules were size-separated by passing the biomass through stainless steel sieves.

### Bioreactor design and operation

Twelve, identical laboratory-scale (2L) glass, EGSB bioreactors were constructed, and operated in four sets of triplicates: the first set (R_S1_–R_S3_) containing only S granules (0.6–1.0 mm); the second set (R_M1_–R_M3_) containing only M-sized granules (1.0–1.4 mm); the third set (R_L1_–R_L3_) containing only L granules (1.4–1.8 mm); and the fourth set (R_N1_–R_N3_) started with the unfractionated, naturally distributed (N) sludge (Fig 5).

Apart from granule size in the starter biomass, the 12 bioreactors, each inoculated with15 gVS L_bioreactor_^−1^, were operated identically for 51 days. The biomass was allowed a 48-h acclimatisation period at 37ºC, regulated using built-in water jackets and recirculating water baths (Grant Optima, T100-ST12), before feeding and recirculation were commenced, which were controlled using peristaltic pumps (Watson and Marlow 2058 and 300 series, respectively). Influent was introduced at the base of each bioreactor, and bioreactor liquor was recirculated through the system to achieve the superficial upflow velocity required (Table 1), according to the same set-up, and approach, as described previously (38, 39).

The saccharide-rich, synthetic feed (Table 2), based on recommendations (40), and supplemented with trace elements (41), was prepared freshly every other day and fed to the 12 bioreactors from a single, thoroughly mixed reservoir to ensure homogeneity.

**Table 2.** Composition of synthetic feed

| **Component** | **Concentration (g L<sup>-1</sup>)** |
| --- | --- |
| Glucose | 3.75 |
| Fructose | 3.75 |
| Sucrose | 3.56 |
| Yeast Extract | 1.45 |
| Urea | 2.15 |
| Sodium Bicarbonate | 10.0* |

\*After Day 6

Sodium bicarbonate was added to the influent on day 6, and for the remainder of the experiment to act as a pH buffer, as the pH of the bioreactor liquor had dropped to 4 during the first week (Phase 1). Some biomass washout was observed over the final two weeks of the trial. Upon take-down, on day 51, biomass was re-fractionated to determine the distribution of granule sizes, and stored for DNA extractions and sequencing.

### Sampling and analytical techniques to monitor bioreactor performance

Biogas concentrations of methane, and effluent concentrations of total COD (tCOD), soluble COD (sCOD), volatile fatty acids (VFA) and pH, were monitored three times a week throughout the 51-d trial. Biogas methane concentrations were determined using a VARIAN CP-3800 gas chromatograph (Varian, Inc., Walnut Creek, CA). pH was measured using a benchtop meter (Hanna Instruments, Woonsocket, RI). COD was measured using pre-prepared COD test kits (Reagacon, Shannon, Ireland) and following the recommendation of the manufacturer. Samples for tCOD assays were each prepared by adding an homogenous sample directly to the test kit, whilst for sCOD, the sample was first centrifuged for 10 min at 14,000 rpm and the supernatant was added to the test kit. COD tests were incubated for 2 h at 150ºC and concentrations were determined using a spectrophotometer (Hach Dr/4000) at 435 nm. VFA contents of supernatant from effluent samples were separated, and quantified, using gas chromatography (Varian 450-GC).

### DNA extraction

For each sample investigated, a mass of 0.1 g wet sludge was transferred to respective, sterile tubes in triplicate. DNA was extracted on ice following the DNA/RNA co-extraction method (42), which is based on bead beating in 5% (w/v) cetyl trimethylammonium bromide (CTAB) extraction buffer, followed by phenol-chloroform extraction. Integrity of nucleic acids was assessed using a NanoDrop^TM^ spectrophotometer (Thermo Fisher Scientific, Waltham, MA, USA), and concentrations were determined using a Qubit fluorometer (Invitrogen, Carlsbad, CA, USA) and normalised to 5 ng DNA µl^−1^ for storage at −80ºC.

### High-throughput DNA sequencing

Partial 16S rRNA gene sequences were amplified using the universal bacterial and archaeal primers, 515F and 806R (43), as previously described (13), and with amplicon sequencing on an Illumina MiSeq platform (at FISABIO, Valencia, Spain).

### Bioinformatics and statistical analysis

Abundance tables were generated by constructing OTUs (as a proxy for species). Statistical analyses were performed in R using the combined data generated from the bioinformatics as well as meta data associated with the study. An OTU table was generated for this study by matching the original barcoded reads against clean OTUs (a total of 2,793 OTUs for *n* = 49 samples) at 97% similarity (a proxy for species-level separation). Alpha diversity analyses included the calculation of Shannon entropies and rarefied richness. Further multivariate integration (MINT) algorithms identified study-wise discriminants with additional detail available in Supplemental Material.

### Data Availability

The sequencing data from this study are available on the European Nucleotide Archive under the study accession number PRJEB28212 (http://www.ebi.ac.uk/ena/data/view/PRJEB28212).

## Supplementary Information

Supplementary information has been uploaded in a separate document for review.

## Acknowledgements

The authors thank NVP Energy for providing anaerobic sludge granules. SC was supported by the Engineering and Physical Sciences Research Council, UK (EP/J00538X/1). CQ was funded by an MRC fellowship MR/M50161X/1 as part of the CLoud Infrastructure for Microbial Genomics (CLIMB) consortium MR/L015080/1. CM was supported by Erasmus and by the University of Turin and NUI Galway. UZI was funded by NERC IRF NE/L011956/1. GC, SM and ACT were supported by a European Research Council Starting Grant (3C-BIOTECH 261330) and by a Science Foundation Ireland Career Development Award (17/CDA/4658) to GC. ACT was further supported by a Thomas Crawford Hayes bursary from NUI Galway, and a Short-Term Scientific Mission grant through the EU COST Action 1302.

## Author Contributions

ACT, SC, UZI and GC designed the study. ACT performed all of the physico-chemical characterisation with assistance from CM, SM, EG, CS, SD and CN. ACT prepared the sequencing libraries. UZI wrote the scripts for data analysis, which was conducted by ACT. Results were interpreted by ACT, CQ, UZI and GC. ACT drafted the paper and UZI and GC revised the document. UZI and GC are joint corresponding authors. All authors approve the paper and agree for accountability of the work therein.

---

**Figure S1.** Bar plot of the biomass yield, calculated on Day 51, for each bioreactor (asterisk indicates R_N2_ bioreactor, which was stopped on day 22).

**Figure S2.** MINT study-wise discriminant analysis where **(a)** shows the first two components of samples (MINT PLS-DA) using all the OTUs with ellipse representing 95% confidence interval and percentage variations explained by these components in axes labels; **(b)** shows the optimum number of discriminating OTUs found for these 2 components identified as diamonds; and **(c)** is similar to **(a)** but the samples are drawn only using the 38 discriminant OTUs (MINT sPLS-DA); (**d – g**) show the MINT sPLS loading vectors *a*_1_ and *a*_2_ with non-zero weights for component 1 and component 2 where **(d)** shows contributions by emerging granules from R_S_; **(e)** from R_M_; **(f)** from R_L_ and **(g)** from R_N_ studies. Loading vectors are coloured by emerging size with maximal abundance (note: while interpreting this figure, focus should be on the colour of the bars and not the positive/negative projections); **(h)** shows the phylogenetic tree (extracted from the global tree) of the discriminant OTUs; **(i)** indicates which MINT sPLS component each OTU is extracted from; **(j)** the heatmap with mean relative abundance values (drawn using EvolView http://www.evolgenius.info/evolview/); and **(k)** the taxonomic classification of discriminant OTUs coloured by unique phyla to which they belong.

**Figure S3.** Top 25 most abundant taxa from the emerging sizes, ordered by size and based up on variances in the 16S rRNA gene

